# Engineering species-like barriers to sexual reproduction

**DOI:** 10.1101/079095

**Authors:** Maciej Maselko, Stephen C. Heinsch, Jeremy Chacón, William Harcombe, Michael J. Smanski

## Abstract

We introduce a novel approach to engineer a genetic barrier to sexual reproduction between otherwise compatible populations. Programmable transcription factors drive lethal gene expression in hybrid offspring following undesired mating events. As a proof of concept, we target the *ACT1* promoter of the model organism *Saccharomyces cerevisiae* using a dCas9-based transcriptional activator. Lethal over-expression of actin results from mating this engineered strain with a strain containing the wild-type *ACT1* promoter.

---

Controlling the exchange of genetic information between sexually-reproducing populations has applications in agriculture, eradication of disease vectors, control of invasive species, and the safe study of emerging biotechnology applications. Specific examples include preventing herbicide resistance genes from moving from cultivated to weedy plant varieties^1^, generating novel mating incompatibilities to control pest populations, and for the safe study of gene-drives^2^. These applications could be achieved by engineering a speciation event, with speciation defined as reproductive isolating mechanisms that prevent genetic exchange between newly formed taxa ^3,4^.

Ideally, the introduction of species-like barriers would result in an engineered organism that behaves and can be propagated in an identical fashion to its non-modified counterpart. Changing the genetic code has been proposed as a means to accomplish this. Genetic recoding has been successful in *Escherichia coli*^5,6^ and *Saccharomyces cerevisiae* may soon follow^7,8^. However, we are not likely to recode higher organisms with ease in the near future. There are a handful of other examples of engineering genetic incompatibility in the literature. A “synthetic species” of *Drosophila melanogaster*^9^ was developed by knocking out the *glass* transcription factor, and integrating a *glass* dependent killing module which is activated when mated with wild-type flies. Some plants may be engineered to only self-fertilize by engineering flowers which never open^10^. However, these are either only applicable to a small number of species and/or dramatically change the engineered organism’s phenotype.

Here we describe a novel and broadly applicable approach to engineer species-like barriers to sexual reproduction. This method interrupts sexual reproduction between populations of different genotypes with minimal effects on growth and reproduction. Further, propagation of the engineered organisms does not require the use of exogenous inputs^11^. This technology may enable more scalable means for the containment of transgenic organisms and provide additional tools to disrupt pest species reproduction.

In our approach, synthetic species-like genetic barriers are introduced via a relatively simple system (Fig. 1a–c) that utilizes programmable transcriptional activator(s) capable of lethal overexpression of endogenous genes. Lethality in the engineered strain is prevented by refactoring the target locus, allowing the programmable activator to be expressed in the engineered strain. This activator serves as a sentinel for undesired mating events. Hybridization between the synthetically incompatible (SI) strain and an organism containing the transcriptional activator’s target sequence results in lethal gene expression (Figure 1b). Because programmable transcription activators have been shown to work in many organisms^12–17^, this technology is expected to readily transfer to higher organisms including plants^18^, insects^12^, and vertebrates^19^ (Figure 1c).

**Figure 1.**
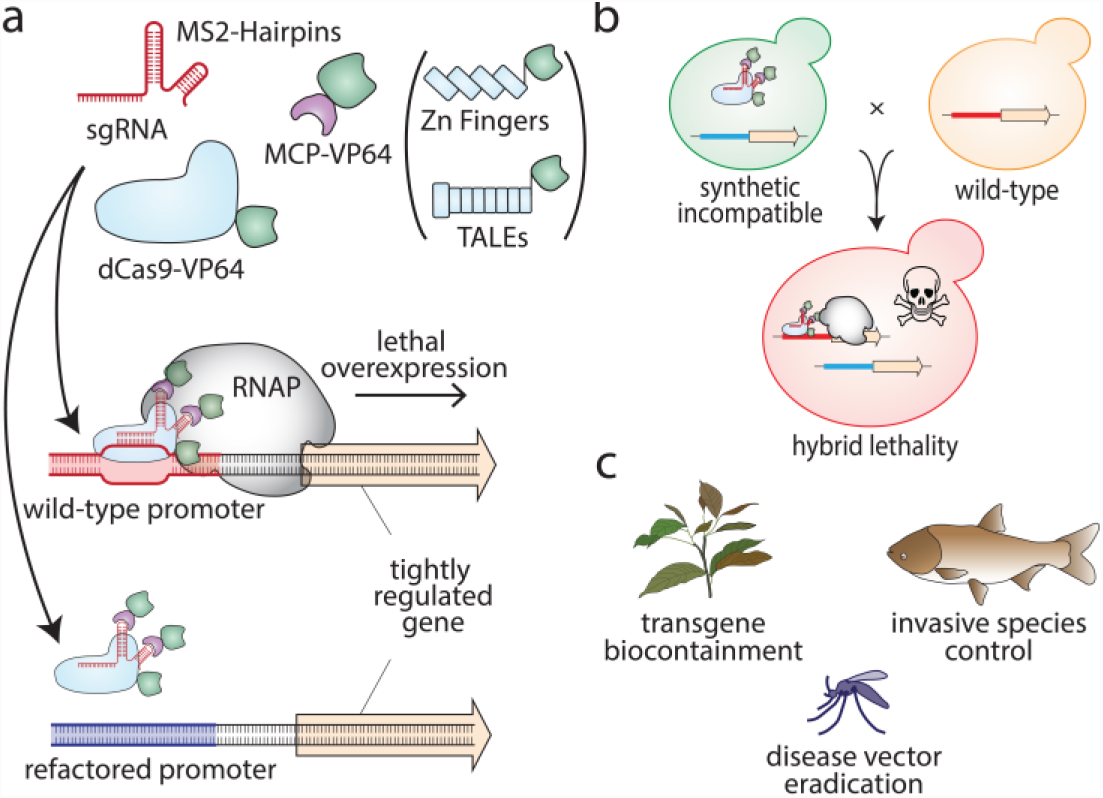
Overview of synthetic incompatibility. **(a)** Macromolecular components that constitute programmable transcription factors (above), and schematic illustration showing lethal gene expression from a wild-type but not a refactored promoter (below) **(b)** Illustration of hybrid lethality upon mating of wild-type (orange cell) and SI (green cell) parents. Macromolecular components are labeled in (a), red DNA signifies WT promoter, and blue DNA signifies refactored promoter. Skull and crossbones indicates a non-viable genotype. **(c)** Possible applications for engineered speciation.

We demonstrate our approach in *S. cerevisiae* using a programmable transcriptional activation system composed of dCas9-VP64 combined with sgRNA aptamer binding MS2-VP64 (referred hereafter as DVM) which is based on previously demonstrated strong activators^17,20^. Appropriate target genes were identified empirically by using the DVM system to activate promoters of genes whose overexpression is reported to generate an ‘inviable’ phenotype in the Saccharomyces Genome Database^21^ (Supplementary Table 1). We designed sgRNAs to bind unique sequences immediately upstream of NGG protospacer adjacent motif (PAM) sites in an approximately 200bp window upstream of predicted transcriptional start sites^22^ of candidate genes. Transformant growth rates were then measured for ~10.5 days. We identified several target sites which generated severely reduced growth rates (Figure 2a, Supplementary Figure 1). A target site on the bottom strand 190 nucleotides upstream of the *ACT1* transcriptional start site resulted in the strongest growth defect with no visible growth after 10 days. The nine PAM distal nucleotides are predicted to be Forkhead transcription factor binding sites^23^.

**Figure 2.**
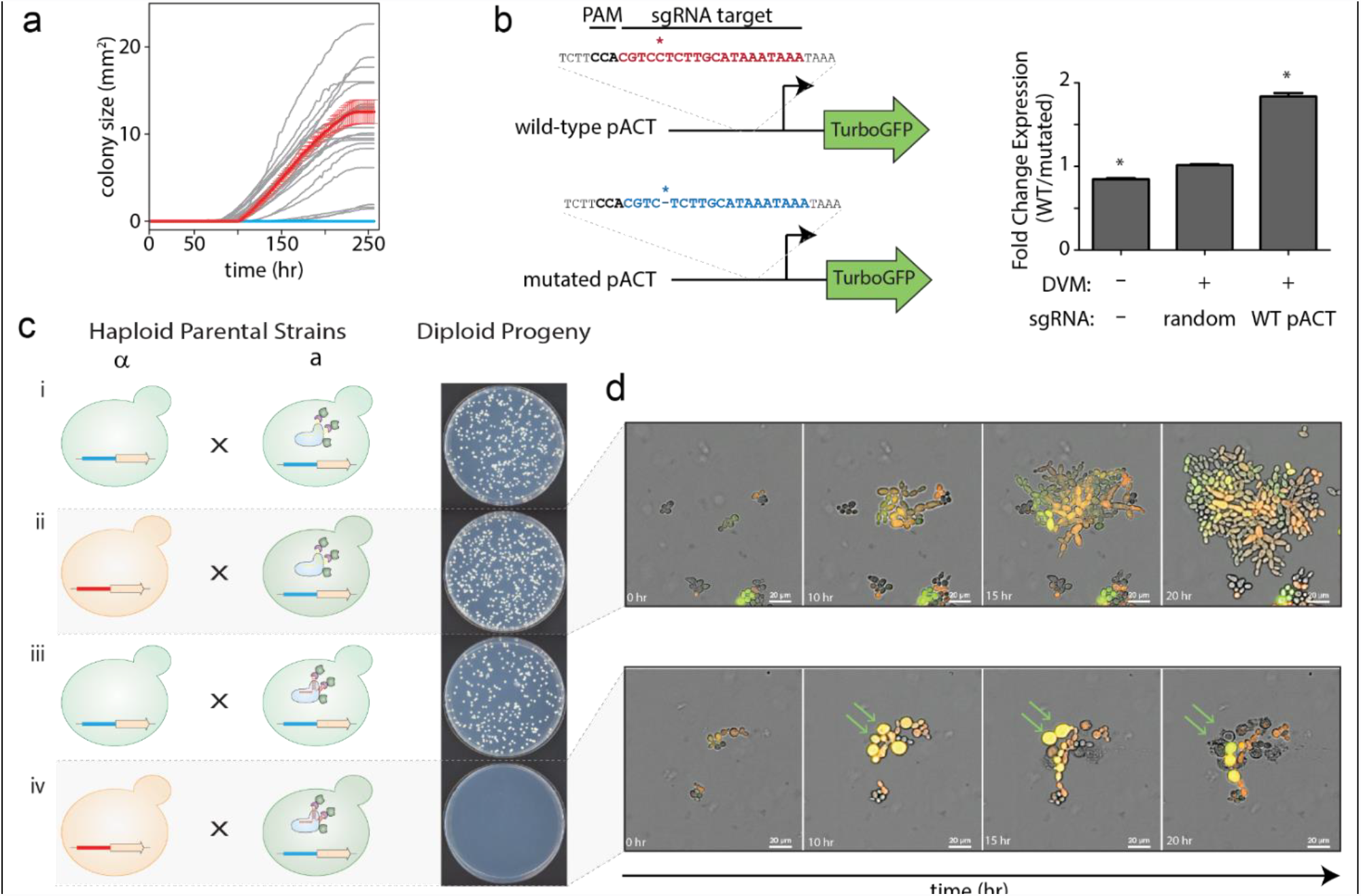
Engineering speciation by synthetic incompatibility. **(a)** Growth curves of yeast expressing DVM targeted to promoter regions of SI candidate genes. Random sgRNA control shown in red. Best *ACT1* targeting sgRNA in light blue. All others in grey. (n=2, +/− SD, error bars omitted for grey lines for clarity) **(b)** (Left) Diagram of mutated and wild-type *ACT1* promoter-GFP constructs. (Right) GFP expression ratios with or without DVM and/or *ACT1* promoter specific sgRNA. (n=3, +/− SD). **(c)** (Left) Schematic representation of SI components present in haploid strain crosses and (Right) the resulting diploid colonies. **(d)** Live cell imaging time lapse of diploid cells from crossing RFP^+^ *MATα* with GFP^+^ *Mata* cells in a compatible (Top) and incompatible (Bottom) mating. Green arrows indicate cells which swell and lyse.

To generate a SI strain, we used Cas9^24^ to introduce a mutation by non-homologous end joining in the *ACT1* promoter. The mutated promoter differs from wild-type by a single cytosine deletion 3 bp upstream of the PAM site. There is no observable growth phenotype resulting from the mutated *ACT1* promoter (Supplemental Figure 2 and Supplementary Note 2). We characterized transcription from the mutated promoter by expressing TurboGFP^25^ under the control of the wild-type or mutated *ACT1* promoters in the presence and absence of DVM (Fig. 2b and Supplemental Figure 3). There was a slight increase in TurboGFP expression from the mutated promoter in the absence of DVM. However, no change was found with a non-targeting sgRNA. TurboGFP expression was 1.8 fold higher from the wild-type *ACT1* promoter than the mutated promoter when DVM was guided by an sgRNA targeting the wild-type promoter. Together, these results indicate that the mutation in the *ACT1* promoter does not substantially change native expression but prevents targeted transcriptional activation by DVM guided to the wild-type sequence. We completed construction of the SI strain by chromosomally integrating a DVM targeted to the wild-type *ACT1* promoter sequence in the strain containing the mutated *ACT1* promoter (*i.e.* Figure 1b).

Next, we examined the genetic compatibility between the SI strain and a strain with the wild-type *ACT1* promoter. *S. cerevisiae* has haploid mating types *MAT*a and *MAT*α, and can be propagated as a haploid of either mating type or as a diploid after mating. We mated haploid strains with different auxotrophic markers and selected for diploids to determine mating efficiency (Figure 2c). Mating a *MAT*a strain with the SI genotype but a random sequence sgRNA to a *MAT*α strain also containing the mutated *ACT1* promoter resulted in numerous diploid colonies (Figure 2c i). This shows that expression of the DVM machinery or mutation of the *ACT1* promoter do not prevent sexual reproduction. This same *MAT*a strain was also successfully mated to a *MAT*α strain carrying the wild-type *ACT1* promoter (Figure 2c ii), as the random sequence sgRNA does not induce lethal overexpression of *ACT1*. We were also able to cross the *MAT*a strain with a complete SI genotype to a *MAT*α strain with the mutated *ACT1* promoter (Figure 2c iii). However, when the SI *MAT*a strain was mated with a *MAT*α strain with wild-type *ACT1* promoter, diploid colonies were seen only in low frequencies (Figure 2c iv). This failed mating reflects the genetic incompatibility of the SI genotype with wild-type.

In order to understand the engineered genetic incompatibility on a cellular level, we performed mating experiments and monitored diploid cells using live cell imaging (Figure 2d, Supplementary Movie 1, and Supplementary Note 1). Diploid yeast resulting from a permissive mating (*e.g.* Figure 2c ii) are able to proliferate and produce a microcolony after 20 hours (Figure 2d top). Diploids arising from the non-permissive mating of wild-type *ACT1* promoter yeast with the SI strain undergo a limited number of divisions before swelling and eventually lysing (Figure 2d bottom). These results are consistent with what we expect from uncontrolled cytoskeletal growth. However, the ability for these yeast to divide a few times before lysis may also provide opportunities for recombination and escape.

Next, we turned our attention to the colonies occasionally found when mating the SI strain to the wild-type. These colonies appeared at frequency of 4.83 × 10^−3^ compared to mating with a compatible strain. Sanger sequencing the *ACT1* promotor found that most (3/5) originated from a cell homozygous for the mutated version of the promoter, suggesting that recombination had taken place between homologous chromosomes. Richardson *et al.* reported single-strand oligonucleotide homology directed repair frequencies of 7 × 10^−3^ in the presence of dCas9 in mammalian cell culture^26^. Therefore, a similar mechanism may be responsible for the apparent mitotic gene conversion observed here. A fourth colony had a mutated MS2-VP64 activator and we were unable to locate mutations in the remaining colony.

In conclusion, we have presented the proof of concept for a novel approach to introducing defined genetic barriers to sexual reproduction. Synthetic incompatibility requires a single, phenotypically-neutral genomic edit and the expression of a transcriptional activator targeting the unedited locus. Recently developed CRISPR/Cas9 based technologies should make it possible to apply synthetic incompatibility in a broad variety of sexually reproductive organisms. Recombination events between the target and mutant loci likely triggered by dCas9 binding indicate that applying this technology in higher organisms will require expressing dCas9 activators only in multicellular stages of life so that it is unlikely that enough cells undergo recombination to rescue the whole organism. Applying synthetic incompatibility to crops engineered to make biofuels or pharmaceuticals may allow for broader cultivation while preventing transgene flow to wild relatives or varieties used for human consumption. Synthetic incompatibility may also find applications in biocontrol of pest organisms by releasing SI males to reduce the fecundity of wild populations.

## Methods

### Plasmids

Plasmid sequences can be found in Supplementary File 1 and primer sequences in Supplementary File 2. Plasmid maps are found in Supplementary Figure 4 and descriptions in Supplementary Table 3.

### Strains and Media

Detailed information for all yeast strains can be found in Supplementary Table 4, Supplementary Figure 5, and Supplementary Note 3. Yeast transformations were performed using the Lithium-acetate method^27^. Chemically competent *E. coli* STBL3 (Thermo Fisher) was used for all plasmid cloning and propagation in LB media (MP) supplemented with appropriate antibiotics. All yeast strains were in the CEN.PK *MAT*a or *MAT*α^28^ background which were a gift from Dr. Claudia Schmidt-Dannert. Yeast were grown at 28-30°C on plates or in liquid culture with 250 rpm agitation. Yeast were cultured in YPD (10 g/L yeast extract, 20 g/L peptone, 20 g/L dextrose), 2X YPD, or synthetic dropout (SD) media (1.7 g/L yeast nitrogenous base, 5 g/L ammonium sulfate, yeast synthetic dropout media supplements (Sigma), 20 g/L dextrose). G418 sulfate resistant yeast were selected on YPD agar with 400 ug/ml G418 Sulfate. Counterselection for KlURA3 was performed using 1 g/L 5-Floroorotic acid.

### Screening Candidate Genes

Screening target genes was performed by transforming yeast strain YMM124 with pMM2-20-1 backbone vectors expressing sgRNA to candidate genes (Supplementary Table 4 and Supplementary Table 3). Transformations were plated onto SD-Ura in 6-well plates and incubated at 30°C. To calculate growth rates of colonies on petri dishes^29,30^, we scanned colonies as they grew using Epson Perfection V19 scanners in two hour intervals for 256 hours. We used image analysis to track the areas of colonies as they grew. This entailed converting RGB scans into HSV colorspace, selecting the V channel, performing a background subtraction, smoothing, and using a threshold to identify biomass. The V channel was selected because it had the highest contrast with the background. The background was the first image in a time-lapse, before any colonies appeared. We smoothed images twice with a fine-grain Gaussian filter (sd = 1 pixel, filter width = 7 pixels) to remove noise. We used a single threshold for all images for consistency. Colony centers were identified by applying regional peak detection to a z-projection through time using the thresholded images. When colonies merged, we used these peaks to find the dividing line between colonies: the peaks were used as seeds in a watershed on a distance-transformed image. Once colony boundaries were identified, the number of “on” pixels within a boundary at each moment in time was counted as the colony’s area. We did not include in the analysis colonies which fell along the edge of the petri dish, which merged with colonies along the edge, or which had an ambiguous number of peaks within a large merged region. To calculate growth rates, we log-transformed the area-over-time data and fit a line in a 12 hour moving window. The maximum slope in each time series was recorded as that colony’s growth rate. The growth rates were analyzed by one-way ANOVA followed by Bonferroni’s post-test comparing each condition to the random sgRNA control.

### Plate Based Mate Assay

Haploid *MATa* yeast strain YMM134 and YMM155 were mated to *MATα* strains YMM125 and YMM141 by combining overnight cultures in YPD to an OD_600_ of 0.1 each in 1 ml YPD. The cultures were then incubated at 30°C for four hours, washed once with water and 30 uL were plated onto SD-Ura/Leu dropout media.

### Flow Cytometry

Flow cytometry was performed using yeast strains YMM158 through YMM163. YMM158, YMM160, and YMM162 expressed TurboGFP driven by the wild-type *ACT1* promoter from plasmid pMM2-17-1. YMM159, YMM161, and YMM163 contained pMM2-17-2 and expressed TurboGPF from a mutated *ACT1* promoter. Overnight cultures grown in 2 mL SD-Complete media were diluted to an OD_600_ = 0.5 and grown for an additional four hours. Cells were collected by centrifugation, washed with DPBS, resuspended in DPBS and placed on ice protected from light prior to analysis. Flow cytometry was performed using a LSRFortessa H0081 cytometer. At least 30,000 TurboGFP positive singlet events were collected per sample. The geometric means of GFP fluorescence intensity were compared using one-way ANOVA followed by Tukey’s post-test for pairwise comparisons.

### Live-Cell Imaging

Yeast strain YMM139 was mated separately with YMM156 and YMM157 in SD-Trp dropout media for 2 hours, pelleted, and resuspended in SD-Ura/Leu/Trp. Mated yeast were loaded onto a CellASIC ONIX diploid yeast plate and supplemented with SD-Ura/Leu/Trp. Cells were imaged using a Nikon Ti-E Deconvolution Microscope System every 6 minutes for 20 hours.

## Acknowledgements

M.M. and M.J.S. are supported by a grant from the University of Minnesota Office of the Vice President of Research. S.H. and M.J.S. are supported by an award from the Damon Runyon Cancer Research Foundation. JC is supported by a grant from the Biocatalysis Initiative at the University of Minnesota.

## Author Contributions

M.M. and M.J.S. conceived the experiments and wrote the manuscript. M.M. performed the strain engineering, mating experiments, and flow cytometry. S.C.H. performed the plate-based growth assays. J.C. and W.H. analyzed plate-based growth assay data. M.M., S.C.H., and M.J.S. conceived and performed all other data analysis.

